# LargeQvalue: A Program for Calculating FDR Estimates with Large Datasets

**DOI:** 10.1101/010074

**Authors:** Andrew Anand Brown

**Affiliations:** Wellcome Trust Sanger Institute, Hinxton, Cambridge, CB10 1HH, UK; NORMENT, KG Jebsen Centre for Psychosis Research, Institute of Clinical Medicine, University of Oslo, Norway

**Keywords:** Multiple testing, false discovery rate, q values, large datasets

## Abstract

This is an implementation of the R statistical software qvalue package [Dabney et al., 2014], designed for use with large datasets where memory or computation time is limiting. In addition to estimating p values adjusted for multiple testing, the software outputs a script which can be pasted into R to produce diagnostic plots and report parameter estimates. This program runs almost 30 times faster and requests substantially less memory than the qvalue package when analysing 10 million p values on a high performance cluster. The software has been used to control for the multiple testing of 390 million tests when analysing a full cis scan of RNA-seq exon level gene expression from the Eurobats project [Brown et al., 2014]. The source code and links to executable files for linux and Mac OSX can be found here: https://github.com/abrown25/qvalue. Help for the package can be found by running `./largeQvalue --help`.

## 1. INTRODUCTION

Recent developments have allowed us to investigate the action of DNA in a new, hypothesis free framework. The ability to collect information on DNA at millions of loci has led to a reappraisal of statistical methods to correct for multiple testing: it has been argued that in many contexts traditional measures of significance such as Bonferroni are too conservative, leading to an unacceptably high rate of falsely accepting null hypotheses. The false discovery rate is an alternative way of accounting for multiple testing, which tolerates a higher false positive rate in order not to discard potentially interesting findings [Benjamini and Hochberg, 1995]. It can simplistically be viewed as the proportion of falsely rejected null hypotheses, and the concept was further developed in Storey and Tibshirani [2003] and Storey et al. [2004], and implemented in the R package qvalue [Dabney et al., 2014].

Advances in sequencing have accelerated our ability to perform experiments of even greater scope. Whereas genotyping arrays ascertain DNA at around a million SNPs, the 1000 Genomes project [1000 Genomes Project Consortium, 2012] has uncovered around 100 million variants in the human genome. Application of sequencing to produce molecular phenotypes has also revolutionised how we can interrogate cellular function with few preconceived notions; for example sequencing of RNA allows us to construct hundreds of thousands of phenotypes, each capturing the expression of a given exon [Lappalainen et al., 2013]. This permits a full, hypothesis free investigation of the genetics of gene regulation. Together, these developments imply that a full genome-wide scan for all possible variants affecting exon expression would involve *∼* 10^13^ hypothesis tests.

The qvalue package [Dabney et al., 2014], implemented in R [R Core Team, 2013], allows estimation of false discovery rates, as well as the quantity π_0_, which is an estimate of the proportion of the tests for which the null hypothesis is true. However, R programs usually run slower and consume more memory than programs written in lower level languages, rendering the qvalue package impractical when the number of tests is of the order 10^13^. Here I present an implementation of qvalue in a compiled language, which runs faster, uses less memory, and is capable of scaling up to the experiments produced by these new sequencing technologies.

## 2. METHODS

### 2.1 Brief description of qvalue algorithms

The qvalue package implements the algorithms described in Storey [2002], Storey and Tibshirani [2003] and Storey et al. [2004]. It takes as input a set of p values and maps these onto q values, which can be interpreted as follows: a decision rule which rejects all null hypotheses with corresponding q values less than *α* has an estimated false discovery rate of *α*. Taking an increasing set of p values *p*_1_*..p_n_* this mapping is defined as:

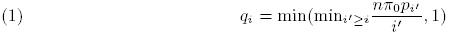

The quantity π_0_ is the proportion of null hypotheses amongst all tests, and is a central difference between the methods used to control the FDR in Benjamini and Hochberg [1995] and the qvalue approach. Its estimation is described in Sections 2.4, 2.5 and 2.6.

### 2.2 Implementation

largeQvalue is written mainly in the D programming language [Alexandrescu, 2010]. D is a compiled language which promises runtime speeds of the same order as C or C++ while guaranteeing memory safety, and also allows easy integration with code written in C. Leveraging this, parts of the program are written in C; in particular we use GNU Software Library (http://www.gnu.org/software/gsl/) functions for sampling from the binomial distribution, and code from the R project [R Core Team, 2013] for fitting smoothing splines. The program is released as a standalone binary for use on linux systems, as well as source code which can be downloaded from a github repository (https://github.com/abrown25/qvalue) and compiled on Windows and Mac OSX platforms. This source code is released under the GNU General Public Licence v3 (http://www.gnu.org/copyleft/gpl.html). The program is called from the command line with options set using parameter flags. A description of all the available options can be found by running `largeQvalue --help`.

### 2.3 Input and output file formats and commands

The input format for this program is a whitespace delimited (i.e. either tab or space separated fields) text file where the p values to be adjusted for multiple testing lie within one column. Consecutive runs of whitespace are collapsed to one whitespace character. The file can either be specified using the `--input` flag (i.e. `--input PValueFile`) or if not present, then following UNIX convention the filename is taken as the last argument on the command line after all arguments have been passed. If no arguments remain, then input is taken from the stdin. The column containing the p values can be specified with the `--col` flag; by default it is assumed to be the first column. If the `--header` flag is not present, then it is assumed the file does not have a header line. Any data that cannot be converted to numerical form will be treated as missing (the corresponding q value estimate will be written as NA). If any numerical values lie outside the interval [0, 1], then the program will terminate with the message `Some p values are outside [0, 1] interval`.

The standard output file is the input file with an additional column on the end which specifies the q value [Storey, 2002] corresponding to the p value in that row. If a filename for the output is not specified using the `--out` flag, then output is written to stdout. The `--sep` flag can be used to specify the character which separates the q value column from all other columns (this option is available to keep consistency with the rest of the file). This can be specified as either `--sep space` (equivalently `--sep s`), or `--sep tab` (`--sep t`). The default option is to use tabs; tabs are also used if the `--sep` flag is mispecified.

An optional parameter file is written, if specified using the `--param` flag. This file contains the estimate of π_0_ used to calculate q values as well as other intermediate parameters. This file can also be run in R, either by pasting it at the R prompt or by running the command `source(“ParameterFileName”)` from the R terminal. This script will produce diagnostic plots, as seen in Figure 1 and described in Sections 2.5 and 2.6. For this functionality, the graphing package ggplot2 [Wickham, 2009] is required.

**Figure 1.**
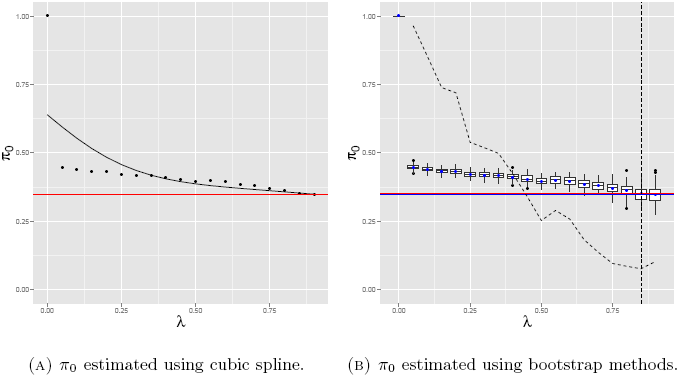
(a) Example of the diagnostic plot produced by the standard options, the dots represent the estimate of the proportion of null hypotheses (π_0_) for various values of λ. These values should converge as λ approaches 1; the point of convergence is estimated using a spline function (black line). Estimate of convergence is at the red line. (b) Example of the diagnostic plot with the `--boot` option. Again, estimates of 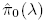 are shown (blue dots). The uncertainty of these estimates is calculated by taking 100 bootstrap samples of the data, the distribution of these parameter estimates is shown by the box plots. The estimate of π_0_ used to calculate q values (the red line) is the value that minimises the mean squared error with the lowest estimate of π_0_ calculated on the original data (blue line).

### 2.4 Estimation of π_0_

One method to estimate the proportion of null hypotheses is to choose a value *λ ∈* [0, 1) and assume that every *p > λ* is from the null hypothesis (obviously this assumption is more valid as λ tends to 1). Then, assuming uniform p values generated by the null hypothesis, this proportion can be reflected back across the interval. This corresponds to estimates of π_0_ as a function of λ:

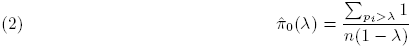

Using the qvalue package in R, the user can specify a value of λ to be used to estimate π_0_ according to Equation 2. The equivalent command in the largeQvalue package uses the flag `--lambda` to specify this value. More sophisticated methods, also implemented in the R package, use a sequence of λ values to produce a range of estimates of π_0_. Information from all of these values is combined either by fitting a cubic spline to these estimates [Storey and Tibshirani, 2003] or by taking bootstrap samples of the p values [Storey et al., 2004]. The implementation of both of these options in largeQvalue is described in the following two sections.

### 2.5 Estimation of π_0_ with cubic splines

The methodology to estimate π_0_ using cubic splines was described in Storey and Tibshirani [2003], and is used when the specified values of λ form a sequence rather than a single number (this is true by default) and the `--boot` flag is not given.

This option takes a sequence of λ values, specified by `--lambda START,STOP,STEP` (default value is 0 to 0.9 in steps of 0.05). Estimates of 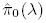 are calculated at every value of λ and a cubic spline is fitted to these points to estimate the asymptote as *λ →* 1. The estimate of π_0_ used is the value of this cubic spline at the maximum value of λ.

The amount of smoothing supplied by the spline can be adjusted using the `--df` flag (default 3, must be between 1 and the length of the λ sequence). If the `--log` flag is specified then the spline is fitted to the 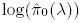 values instead of 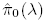.

Figure 1a shows the output of the parameter file processed by R when this methodology is used. The dots show the estimates of π_0_ at various values of λ, the black line is a cubic spline fitted to these points which is used to approximate the function π_0_(λ). The estimate 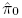 used to estimate q values is shown in red line.

### 2.6 Estimation of π_0_ from bootstrap samples

Storey et al. [2004] proposed a method of estimating π_0_ based on bootstrap samples of the p values. This is used when largeQvalue is run with the flag `--boot`. The seed for the generation of the samples can be set with the flag `--seed` (default 0).

Again, a sequence of λ values is given `--lambda START,STOP,STEP` which is used to calculate estimates 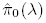. Then, we take 100 bootstrap samples *B_j_* of the p values with replacement and calculate estimates of 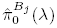 on these new datasets. The implementation of the bootstrap differs from that used in the qvalue package. To save memory and computation time, instead of resampling individual p values we sample counts of p values lying within intervals defined by the sequence of λ from a multinomial distribution. This is effectively the same procedure.

The estimate 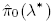 is used to estimate q values, where *λ^*^* is chosen to minimise Equation 3.

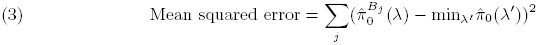

Figure 1b shows the diagnostic plot when this method is applied. Estimates 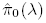 are shown by the blue dots, with 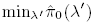 shown by the blue line. The bootstrap estimates 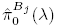 are shown in the box plots and the mean square error by the dashed line. The chosen estimate of π_0_, which minimises this mean squared error, is shown by the red line.

### 2.7 Other parameter flags

The flags `--help` and `--version` will bring up the help page and print the version number of the software respectively. The flag `--issorted` informs the program that the p values are already in ascending order with no missing values: especially with large datasets this can speed up the program considerably.

The flag `--param` allows the user to specify an output file to write estimates of parameters, including the estimate 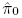 produced by the applied method. When only a single value of λ is specified, this file only contains this point estimate of π_0_. When λ is specified as a sequence, these files also include the estimates 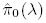 for every value in this sequence. If the cubic spline method is employed, then smoothed cubic splines values are reported at every value of λ. For the bootstrap method the file contains all bootstrap estimates 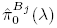 and estimates of the mean square error. Examples of these files can be found in Supplementary Materials S2 and S3. These files can be pasted into R to produce diagnostic plots, examples of which can be seen in Figures 1a and 1b.

The `--pi0` flag allows the estimate of π_0_ to be directly specified.

The `--robust` flag produces q values which are more conservative for very small values of p. Instead of q values defined by Equation 1, we use the formula:

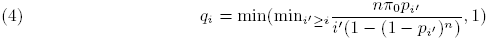

### 2.8 An example command

For the following command: `./largeQvalue --header --col 4 --out OutputFile --param ParameterFile --lambda 0,0.9,0.05 --robust QTLresults.txt`

The p values can be found in the 4th column of QTLresults.txt, write the results to OutputFile and the estimated parameter values to ParameterFile. Use values of λ from 0 to 0.9 in steps of 0.05 to estimate proportion of null hypotheses (standard settings in qvalue) and produce estimates of q values robust for small p values.

### 2.9 Comparison of speed and memory usage of largeQvalue and qvalue package

We compared the CPU time and memory used by largeQvalue with that of the qvalue implementation in R. Each program was run five times against four datasets; these consisted of text files with a single column of either 5 *×* 10^5^, 10^6^, 5 *×* 10^6^, or 10^7^ uniformly distributed numbers between 0 and 1. The following R code was used to call the qvalue algorithm with the cubic spline method:

~~~
library(qvalue)
Pvalues <- scan(file = “InputFile”)
write(qvalue(Pvalues)$qvalues, file = “Qvalues”, ncolumns = 1)
~~~

For the bootstrap method the following code was used:

~~~
library(qvalue)
Pvalues <- scan(file = “InputFile”)
write(qvalue(Pvalues, pi0.method == “bootstrap”)$qvalues,
 file = “Qvalues”, ncolumns = 1)
~~~

The corresponding commands for the largeQvalue package were:

~~~
./largeQvalue --out Qvalues InputFile
~~~

and

~~~
./largeQvalue --boot --out Qvalues InputFile
~~~

### 2.10. List of options

List of options:

Usage: `largeQvalue [options]`

Options:

--help: Print help and quit.
--version: Print version and quit.
--input: File to containing p values to analysis. This can also be specified by the last argument on the command-line after all others have been parsed. If neither are present, it is taken from the stdin.
--out: File to write results to (default = stdout).
--param: Print out parameter list to specified file.
--header: Input has header line (default = FALSE).
--col: Column with p values (default = 1).
--sep: Separator to use to separate the column with q values. Specified as either space or tab (which can be shortened to s or t), (default = tab).
--issorted: File has already been sorted with no missing values (default = FALSE).
--pi0: Use given value of π_0_.
--lambda: Either a fixed number or a sequence given as 0,0.9,0.05 (start,end,step) (default = 0,0.9,0.05).
--robust: More robust values for small p values (default = FALSE).
--df: Number of degrees of freedom used by the spline when estimating pi0 (default = 3).
--log: Smoothing spline applied to log π_0_ values (default = FALSE).
--boot: Apply bootstrap method to find π_0_ (default = FALSE).
--seed: Set seed for generating bootstrap samples (default = 0, equivalent to gsl default).

### 2.11 Binary executables

The binary for 64bit linux can be found here: ftp://ftp.sanger.ac.uk/pub/resources/software/largeqvalue/largeQvalue.tar.gz.

### 2.12 Building from source

To compile an executable binary from source (for example if you are using Windows or Mac OSX), it is necessary to have both a D compiler (we have used the gdc compiler as it produces faster programs) and the gsl library installed on your system. Binaries for the gdc compiler can be downloaded from here: http://gdcproject.org/downloads/ and a link to the gsl software library found here: http://www.gnu.org/software/gsl/. Once both are installed you can clone the github account by running git clone https://github.com/abrown25/qvalue.git. Then change to the qvalue directory and run `make`. Once this is done, if installation has worked then running `./largeQvalue` should bring up the help information. A simple test that the program is working correctly can be run by typing `make test`. If there are any issues with this procedure, please contact.

## 3. RESULTS

### 3.1 CPU time and memory usage of largeQvalue and the qvalue package

To compare performance of the two implementations of the qvalue algorithm, we analysed four different datasets five times with both of the programs on the High Performance Cluster at the Wellcome Trust Sanger Institute (further details in Section 2.9). Table S1 lists the CPU time and maximum memory reported by the cluster. The CPU time is shown in Figure 2. The largeQvalue package requires less time to analyse the data, and the difference increases as the number of tests increases. When there are 10^7^ p values, the largeQvalue program is 27 times faster than the qvalue package when comparing median running times.

We do not believe that the cluster reports reliable measures of memory usage, especially as the largeQvalue package runs for a relatively brief time, and allocates memory for much of it. However, when comparing maximum reported memory use across all runs with 10^7^ p values, this program uses less than a quarter of the memory of the qvalue package.

### 3.2 Application to find tissue specific eQTL

The program was used to look for evidence of cell type specific eQTL in the skin expression data from the Eurobats project [Brown et al., 2014]. We are currently preparing a manuscript based on this work. In brief, a measure of cell type composition was constructed, quantifying the proportions of dermis and epidermis in each individual sample. Then, for every gene and every SNP within 1Mbp of that gene, we tested for interactions between SNP and cell-type measure which affected the expression of that gene (as quantified by RNA-seq), a total of 390,537,631 tests. We ran largeQvalue on these p values. Although the high multiple testing burden meant that we found no significant hits, we estimated a value of π_0_ of 0.961, suggesting that 3.9% of cis-SNPs had different effects on dermis expression in comparison to expression within the epidermis layer. The program took 1 hour 25 mins 55 seconds of CPU time to run, and used 3.99GB of memory.

**Figure 2.**
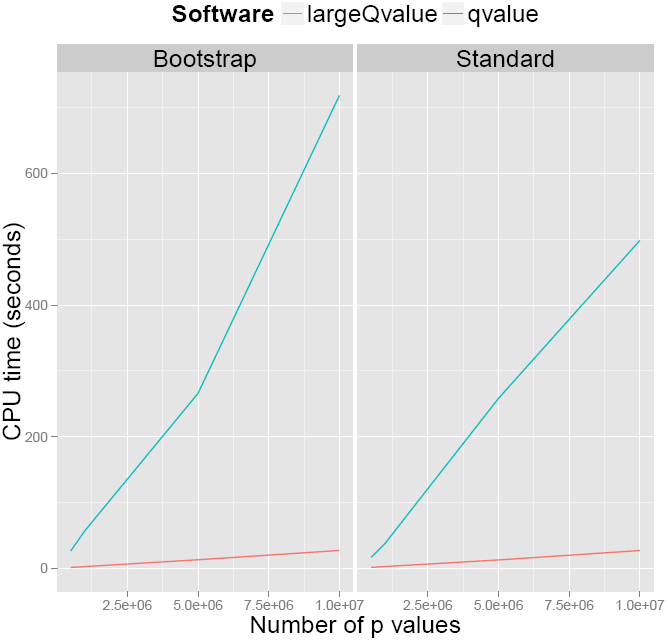
Graph showing amount of CPU time needed to run largeQvalue compared to the qvalue package in R.

## 4 CONCLUSION

The qvalue package has been widely adopted to control for multiple testing; the two papers proposing its methodology [Storey and Tibshirani, 2003, Storey et al., 2004] have been cited over 5,600 times. This means that the procedure and its implementation have become widely accepted. The integration within the R framework makes it easy for results to be processed and plotted. The implementation is capable of analysing the results of genome-wide association studies or studies of differential expression of genes (one context where the methodology was originally proposed). However, it is doubtful that it scales up to cope with the results coming from modern cellular sequencing experiments, which can test hundreds of thousands of phenotypes for association with tens of thousands of SNPs (if concentrating on the cis window). For this we propose our implementation, which has already been used to analyse such data.

## ACKNOWLEDGEMENTS

I would like to thank Leo Parts and Ana Viñuela for their comments and suggestions which have greatly changed this article for the better. Andrew Brown has been supported by a grant from the South-Eastern Norway Health Authority (No. 2011060).

## 1 Supplementary Materials

S1. **CPU and memory usage for largeQvalue and qvalue programs.**

**Table S1.**
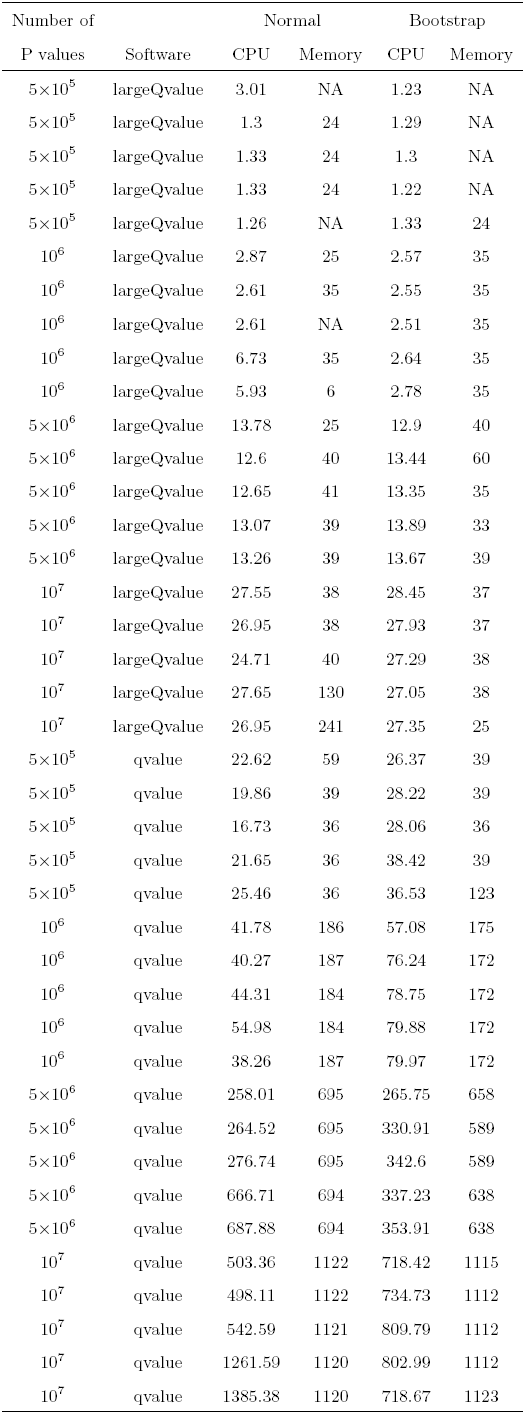
Maximum amount of RAM used, and total CPU time when running largeQvalue and qvalue package to adjust for multiple testing.

| Number of<br>P values | Software | Normal |  | Bootstrap |  |
| --- | --- | --- | --- | --- | --- |
|  |  | CPU | Memory | CPU | Memory |
| $5 \times 10^5$ | largeQvalue | 3.01 | NA | 1.23 | NA |
| $5 \times 10^5$ | largeQvalue | 1.3 | 24 | 1.29 | NA |
| $5 \times 10^5$ | largeQvalue | 1.33 | 24 | 1.3 | NA |
| $5 \times 10^5$ | largeQvalue | 1.33 | 24 | 1.22 | NA |
| $5 \times 10^5$ | largeQvalue | 1.26 | NA | 1.33 | 24 |
| $10^6$ | largeQvalue | 2.87 | 25 | 2.57 | 35 |
| $10^6$ | largeQvalue | 2.61 | 35 | 2.55 | 35 |

Table S1 – *Continued from previous page*
|  | Software | Normal |  | Bootstrap |  |
| --- | --- | --- | --- | --- | --- |
|  |  | CPU | Memory | CPU | Memory |
| $10^6$ | largeQvalue | 2.61 | NA | 2.51 | 35 |
| $10^6$ | largeQvalue | 6.73 | 35 | 2.64 | 35 |
| $10^6$ | largeQvalue | 5.93 | 6 | 2.78 | 35 |
| $5 \times 10^6$ | largeQvalue | 13.78 | 25 | 12.9 | 40 |
| $5 \times 10^6$ | largeQvalue | 12.6 | 40 | 13.44 | 60 |
| $5 \times 10^6$ | largeQvalue | 12.65 | 41 | 13.35 | 35 |
| $5 \times 10^6$ | largeQvalue | 13.07 | 39 | 13.89 | 33 |
| $5 \times 10^6$ | largeQvalue | 13.26 | 39 | 13.67 | 39 |
| $10^7$ | largeQvalue | 27.55 | 38 | 28.45 | 37 |
| $10^7$ | largeQvalue | 26.95 | 38 | 27.93 | 37 |
| $10^7$ | largeQvalue | 24.71 | 40 | 27.29 | 38 |
| $10^7$ | largeQvalue | 27.65 | 130 | 27.05 | 38 |
| $10^7$ | largeQvalue | 26.95 | 241 | 27.35 | 25 |
| $5 \times 10^5$ | qvalue | 22.62 | 59 | 26.37 | 39 |
| $5 \times 10^5$ | qvalue | 19.86 | 39 | 28.22 | 39 |
| $5 \times 10^5$ | qvalue | 16.73 | 36 | 28.06 | 36 |
| $5 \times 10^5$ | qvalue | 21.65 | 36 | 38.42 | 39 |
| $5 \times 10^5$ | qvalue | 25.46 | 36 | 36.53 | 123 |
| $10^6$ | qvalue | 41.78 | 186 | 57.08 | 175 |
| $10^6$ | qvalue | 40.27 | 187 | 76.24 | 172 |
| $10^6$ | qvalue | 44.31 | 184 | 78.75 | 172 |
| $10^6$ | qvalue | 54.98 | 184 | 79.88 | 172 |
| $10^6$ | qvalue | 38.26 | 187 | 79.97 | 172 |
| $5 \times 10^6$ | qvalue | 258.01 | 695 | 265.75 | 658 |
| $5 \times 10^6$ | qvalue | 264.52 | 695 | 330.91 | 589 |
| $5 \times 10^6$ | qvalue | 276.74 | 695 | 342.6 | 589 |
| $5 \times 10^6$ | qvalue | 666.71 | 694 | 337.23 | 638 |
| $5 \times 10^6$ | qvalue | 687.88 | 694 | 353.91 | 638 |
| $10^7$ | qvalue | 503.36 | 1122 | 718.42 | 1115 |

Table S1 – *Continued from previous page*
|  | Software | Normal |  | Bootstrap |  |
| --- | --- | --- | --- | --- | --- |
|  |  | CPU | Memory | CPU | Memory |
| $10^7$ | qvalue | 498.11 | 1122 | 734.73 | 1112 |
| $10^7$ | qvalue | 542.59 | 1121 | 809.79 | 1112 |
| $10^7$ | qvalue | 1261.59 | 1120 | 802.99 | 1112 |
| $10^7$ | qvalue | 1385.38 | 1120 | 718.67 | 1123 |

S2. **Parameter output file using default settings.** This is an example of the standard output using the 

~~~
--param
~~~

 flag and the standard, spline estimation of π_0_. It has been edited to solely to ensure lines fit comfortably on the page and to replace unicode characters.

~~~
#The estimated value of pi0 is: 0.34873
~~~

~~~
#lambda values to calculate this were: [0, 0.05, 0.1, 0.15, 0.2, 0.25, 0.3,
     0.35, 0.4, 0.45, 0.5, 0.55, 0.6, 0.65, 0.7, 0.75, 0.8, 0.85, 0.9]
~~~

~~~
#with the corresponding pi0 values: [1, 0.445501, 0.439427, 0.432638,
    0.431452, 0.419355, 0.418433, 0.415881, 0.409677, 0.403519, 0.394194,
    0.397133, 0.393548, 0.385253, 0.378495, 0.370323, 0.362903, 0.350538,
    0.348387]

#and spline-smoothed pi0 values: [0.639351, 0.594972, 0.553265, 0.515957,
    0.483864, 0.457138, 0.435493, 0.41835, 0.404987, 0.394651, 0.386623,
    0.380249, 0.374948, 0.370262, 0.365875, 0.361593, 0.357319, 0.353028,
    0.34873]

###R code to produce diagnostic plots for spline estimates of pi0:

plot.data <- data.frame(lambda = c(0, 0.05, 0.1, 0.15, 0.2, 0.25, 0.3, 0.35,
    0.4, 0.45, 0.5, 0.55, 0.6, 0.65, 0.7, 0.75, 0.8, 0.85, 0.9),
    pi0 = c(1, 0.445501, 0.439427, 0.432638, 0.431452, 0.419355, 0.418433,
  0.415881, 0.409677, 0.403519, 0.394194, 0.397133, 0.393548,
 0.385253, 0.378495, 0.370323, 0.362903, 0.350538, 0.348387),
pi0Est = c(0.639351, 0.594972, 0.553265, 0.515957, 0.483864, 0.457138,
     0.435493, 0.41835, 0.404987, 0.394651, 0.386623, 0.380249,
     0.374948, 0.370262, 0.365875, 0.361593, 0.357319, 0.353028,
     0.34873))
library(ggplot2)
plot1 <- ggplot(data = plot.data, aes(x = lambda, y = pi0)) + geom_point() +
   geom_line(aes(x = lambda, y = pi0Est)) +
   geom_abline(slope = 0, intercept = plot.data$pi0Est[nrow(plot.data)],
      col = ‘red’) +
   labs(x = expression(lambda), y = expression(pi[0])) +
   theme(axis.title = element_text(size = rel(2))) +
   ylim(0, 1)
print(plot1)
~~~

S3. **Parameter output file using** 

~~~
--boot
~~~

 **flag.** This is an example of the standard output using the 

~~~
--param
~~~

 flag and the 

~~~
--boot
~~~

 option, meaning π_0_ is estimated using bootstrap methods. It has been edited to ensure lines fit comfortably on the page, and to remove many of the 1900 bootstrap π_0_ values outputted as standard (therefore it will not work if directly pasted into R).

~~~
#The estimated value of pi0 is: 0.350538

#lambda values to calculate this were: [0, 0.05, 0.1, 0.15, 0.2, 0.25, 0.3,
  0.35, 0.4, 0.45, 0.5, 0.55, 0.6, 0.65, 0.7, 0.75, 0.8, 0.85, 0.9]
#with the corresponding pi0 values: [1, 0.445501, 0.439427, 0.432638,
  0.431452, 0.419355, 0.418433, 0.415881, 0.409677, 0.403519, 0.394194,
  0.397133, 0.393548, 0.385253, 0.378495, 0.370323, 0.362903, 0.350538,
  0.348387]

#and mean squared error estimates: [42.4599, 0.964491, 0.85278, 0.738637,
   0.719512, 0.538467, 0.518636, 0.497835, 0.423078, 0.340746, 0.252589,
   0.28942, 0.257659, 0.182415, 0.136385, 0.0955588, 0.0851119, 0.0762539,
   0.102196]
###R code to produce diagnostic plots for bootstrap estimates of pi0:

plot.pi0.data <- data.frame(x = rep(c(0, 0.05, 0.1, 0.15, 0.2, 0.25, 0.3,
   0.35, 0.4, 0.45, 0.5, 0.55, 0.6, 0.65, 0.7, 0.75, 0.8, 0.85, 0.9), 100)
   ,
     y = c(1, 0.444482, 0.430108, 0.423909,……
       , 0.393548, 0.392473, 0.385806, 0.375806, 0.356989, 0.341935))
plot.data <- data.frame(x = c(0, 0.05, 0.1, 0.15, 0.2, 0.25, 0.3, 0.35, 0.4,
     0.45, 0.5, 0.55, 0.6, 0.65, 0.7, 0.75, 0.8, 0.85, 0.9),
     y = c(1, 0.445501, 0.439427, 0.432638, 0.431452, 0.419355, 0.418433,
        0.415881, 0.409677, 0.403519, 0.394194, 0.397133, 0.393548,
        0.385253, 0.378495, 0.370323, 0.362903, 0.350538, 0.348387),
mse = c(42.4599, 0.964491, 0.85278, 0.738637, 0.719512, 0.538467,
        0.518636, 0.497835, 0.423078, 0.340746, 0.252589, 0.28942,
        0.257659, 0.182415, 0.136385, 0.0955588, 0.0851119, 0.0762539,
        0.102196),
minpi0 = 0.348387,
final = 0.350538)

library(ggplot2)
plot1 <- ggplot(plot.data, aes(x = x, y = y)) + geom_boxplot(data = plot.pi
  0.data, aes(x = x, y = y, group = x)) +
  geom_point(colour=‘blue’) +
  geom_hline(yintercept = plot.data$minpi0, colour = ‘blue’) +
  geom_line(aes(x = x, y = mse), linetype = ‘dashed’) +
  geom_hline(yintercept = plot.data$final, colour = ‘red’) +
  geom_vline(xintercept = plot.data$x[plot.data$mse==min(plot.data$mse) ], linetype = ‘dashed’) +
  ylim(0,1) +
  labs(x = expression(lambda), y = expression(pi[0])) +
  theme(axis.title = element_text(size = rel(2)))
print(plot1)
~~~

## References

1000 Genomes Project Consortium. An integrated map of genetic variation from 1,092 human genomes. Nature, 491(7422):56–65, 2012.

Andrei Alexandrescu. The D Programming Language. Addison-Wesley Professional, 1st edition, 2010. ISBN 0321635361, 9780321635365.

Yoav Benjamini and Yosef Hochberg. Controlling the false discovery rate: a practical and powerful approach to multiple testing. Journal of the Royal Statistical Society. Series B (Methodological), pages 289–300, 1995.

Andrew Anand Brown, Alfonso Buil, Ana Vinũela, Tuuli Lappalainen, Hou-Feng Zheng, J Brent Richards, Kerrin S Small, Timothy D Spector, Emmanouil T Dermitzakis, and Richard Durbin. Genetic interactions affecting human gene expression identified by variance association mapping. eLife, 3, 2014.

Alan Dabney, John D. Storey, and with assistance from Gregory R. Warnes. qvalue: Q-value estimation for false discovery rate control, 2014. R package version 1.34.0.

Tuuli Lappalainen, Michael Sammeth, Marc R Friedländer, Peter AC’t Hoen, Jean Monlong, Manuel A Rivas, Mar Gonzàlez-Porta, Natalja Kurbatova, Thasso Griebel, Pedro G Ferreira, et al. Transcriptome and genome sequencing uncovers functional variation in humans. Nature, 2013.

R Core Team. R: A Language and Environment for Statistical Computing. R Foundation for Statistical Computing, Vienna, Austria, 2013. URL http://www.R-project.org/.

John D Storey. A direct approach to false discovery rates. Journal of the Royal Statistical Society: Series B (Statistical Methodology), 64(3):479–498, 2002.

John D Storey and Robert Tibshirani. Statistical significance for genomewide studies. Proceedings of the National Academy of Sciences, 100(16):9440–9445, 2003.

John D Storey, Jonathan E Taylor, and David Siegmund. Strong control, conservative point estimation and simultaneous conservative consistency of false discovery rates: a unified approach. Journal of the Royal Statistical Society: Series B (Statistical Methodology), 66(1):187–205, 2004.

Hadley Wickham. ggplot2: elegant graphics for data analysis. Springer New York, 2009. ISBN 978-0-387-98140-6. URL http://had.co.nz/ggplot2/book.

